# ASEP: gene-based detection of allele-specific expression in a population by RNA-seq

**DOI:** 10.1101/798124

**Authors:** Jiaxin Fan, Jian Hu, Chenyi Xue, Hanrui Zhang, Muredach P. Reilly, Rui Xiao, Mingyao Li

## Abstract

Allele-specific expression (ASE) analysis, which quantifies the relative expression of two alleles in a diploid individual, is a powerful tool for identifying *cis*-regulated gene expression variations that underlie phenotypic differences among individuals. Existing methods for gene-level ASE detection analyze one individual at a time, therefore wasting shared information across individuals. Failure to accommodate such shared information not only loses power, but also makes it difficult to interpret results across individuals. However, ASE detection across individuals is challenging because the data often include individuals that are either heterozygous or homozygous for the unobserved *cis*-regulatory SNP, leading to heterogeneity in ASE as only those heterozygous individuals are informative for ASE, whereas those homozygous individuals have balanced expression. To simultaneously model multi-individual information and account for such heterogeneity, we developed ASEP, a mixture model with subject-specific random effect accounting for multi-SNP correlations within the same gene. ASEP is able to detect gene-level ASE under one condition and differential ASE between two conditions (e.g., pre-versus post-treatment). Extensive simulations have demonstrated the convincing performance of ASEP under a wide range of scenarios. We further applied ASEP to RNA-seq data of human macrophages, and identified genes showing evidence of differential ASE pre-versus post-stimulation, which were extended through findings in cardiometabolic trait-relevant genome-wide association studies. To the best of our knowledge, ASEP is the first method for gene-level ASE detection at the population level. With the growing adoption of RNA-seq, we believe ASEP will be well-suited for various ASE studies for human diseases.

## INTRODUCTION

Genome-wide association studies (GWAS) are successful in identifying candidate loci for complex human diseases and traits.^1,2^ Despite the impressive success for disease susceptibility loci discovery, few, if any, results from GWAS have led yet to the delivery of new therapies.^3^ The association peaks from GWAS typically identify a handful of gene candidates, but it is often unclear whether these candidates are expressed in relevant tissues and cell types. Further complicating the picture, we now know that most GWAS signals are probably the result of regulatory variants that impact gene expression, rather than amino acid changes. Data on gene expression from tissues and cell types directly involved in disease are critically important to find causative genes.

A commonly used approach to understand the functional roles of GWAS identified genetic variants is expression quantitative trait loci (eQTL) analysis.^4, 5^ The rationale is that, a genetic variant, known as an eQTL, influences the expression level of a gene, and differences in gene expression levels among individuals may lead to different phenotypes. Studies have found that many GWAS identified single nucleotide polymorphisms (SNPs) are significantly enriched for eQTLs, compared to control SNPs matched by allele frequencies.^6^ eQTL analysis identifies both *cis*- and *trans*-regulatory SNPs, in which *cis*-eQTLs affect gene expression in an allele-specific manner, with implications on underlying mechanism, whereas *trans*-eQTLs affect gene expression in an allele independent manner.^5^ Although eQTL analysis has successfully uncovered functional variant loci that regulate gene expression, typical eQTL analysis only tells *local* versus *distal* association.^7, 8^ The lack of explicit information on *cis*- versus *trans*-makes it difficult to directly link to underlying mechanism, and the requirement of a relatively large sample size for eQTL analysis further makes it impractical for studies that involve difficult-to-collect tissues.^9^

An alternative approach to eQTL is analysis of allele-specific expression (ASE), which refers to unequal expression between paternal and maternal alleles of a gene in a diploid individual, driven by *cis*-regulatory variants located near the gene.^10^ The allelic imbalance of gene expression may explain phenotypic variation and disease pathophysiology. Since the two alleles used to measure ASE are expressed in the same cellular environment and genetic background, they can serve as internal control and eliminate the influence of *trans*-acting genetic and environmental factors. It has been shown that ASE analysis requires 8-fold less samples than eQTL analysis to reach the same power in detecting *cis*-regulatory SNPs, and is less sensitive to SNPs with low minor allele frequencies (MAFs) compared to eQTL analysis.^11^

To understand the role of ASE in human diseases, we exploit allelic imbalance by RNA sequencing (RNA-seq), which provides allele-specific read counts at exonic SNPs distinguished by heterozygous sites.^12^ Existing methods for ASE detection report evidence of ASE in single individuals, in which the ASE is quantified for each SNP (e.g., QuASAR^13^), and a gene-level ASE is obtained by integrating effects across SNPs in the same gene for an individual (e.g., MBASED^14^ and GeneiASE^15^). However, evidence of ASE is often shared across individuals. Failure to accommodate such shared information not only loses power, but also makes it difficult to interpret results across individuals. It is desirable to have a method that simultaneously models both multi-SNP and multi-individual information.

ASE detection across individuals, however, is challenging because the data often include individuals that are either heterozygous or homozygous for the unobserved *cis*-regulatory SNPs, leading to heterogeneity in ASE. Such heterogeneity complicates the analysis because only heterozygous individuals are informative for ASE, whereas those homozygous individuals have balanced expression. Further, when analyzing multiple SNPs in the same gene, haplotype phase information is needed to separate the paternal and maternal alleles. Although it is possible to infer haplotype phase from DNA sequencing, most studies do not have such data available. Even when phase information is available, cross-individual read count alignment is still needed when performing cross-individual analysis. **Figure 1** illustrates these analytical challenges in cross-individual gene-based ASE analysis.

**Figure 1.**
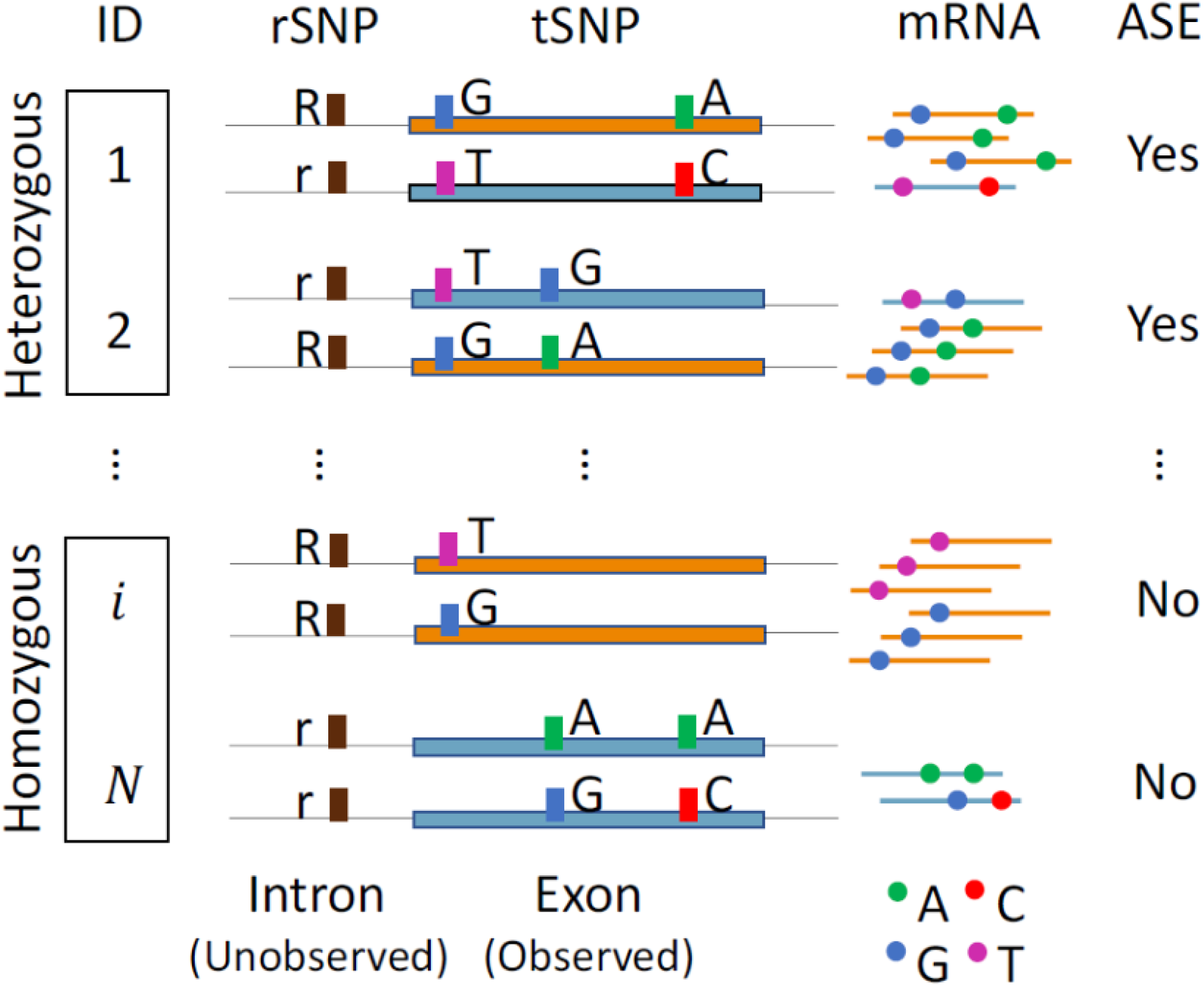
Challenges in cross-individual gene-based ASE analysis. Heterogeneity of the ASE effect exists across individuals in a population. Because the *cis-*regulatory SNP (*rSNP*) is often unobserved, the bulk RNA-seq data include individuals (ID) that are either heterozygous or homozygous at the *rSNP*. The mRNA expression levels differ between two haplotypes only in those heterozygous individuals. Additionally, a gene may have multiple heterozygous transcribed SNPs (*tSNP*s). To differentiate paternal and maternal alleles, haplotype phase information is needed, which is often not available in most studies. Further complicating the analysis, to aggregate ASE effects across individuals, haplotypes that reside on the same allele of the unobserved *rSNP* need to be aligned across individuals.

To properly perform cross-individual gene-based ASE analysis, we propose ASEP (<u>A</u>llele-<u>S</u>pecific <u>E</u>xpression analysis in a <u>P</u>opulation), a mixture model with subject-specific random effect to account for correlation of multiple SNPs within the same gene. ASEP is able to detect gene-level ASE under one condition and differential ASE between two conditions (e.g., pre-versus post-treatment). Through extensive simulations and analysis of a real RNA-seq data set from human transcriptomic studies, we demonstrate that combining shared ASE information across SNPs and individuals leads to easier interpretation and improved power in identifying genes with ASE. Results from our analysis shed light on the functional roles of GWAS identified genetic variants.

## MATERIAL AND METHODS

### Motivation and overview

The primary goal of ASEP is to perform gene-based ASE analysis at the population level. However, the population is a mixture of individuals that are either heterozygous or homozygous at the unobserved *cis*-regulatory SNP. When considering all individuals together, evidence of ASE from heterozygous individuals will be diluted by the inclusion of balanced expression from homozygous individuals. To account for such heterogeneity, we develop ASEP, a generalized linear mixed-effects model in which the subject-specific random effect is assumed to follow a mixture distribution, to detect gene-level ASE under one condition and differential ASE between two conditions in the population.

### Notation

For a given gene *g*, let *rSNP* be its *cis*-regulatory SNP with alleles *R* and *r*, where the *R* allele leads to increased expression level of the residing haplotype as compared to the *r* allele. For individuals with homozygous or heterozygous *rSNP*, we denote them as from ‘Hom’ or ‘Het’ group of individuals, respectively. Let *tSNP* be the heterozygous transcribed SNP. Homozygous transcribed SNPs are excluded from the analysis since they do not provide information on allelic expression.

### Detection of ASE under one condition

For individual *i* at *tSNP j* of gene *g*, let *X*_*ij*_ be the read count for the reference allele in the genome, and *Y*_*ij*_ be the total read count at the SNP. Assume haplotype phase information is known, i.e., the paternal and maternal alleles can be differentiated when there are more than one *tSNP*s of the gene. We further assume the major haplotype, defined as the haplotype that resides with the *R* allele of the *rSNP* which has higher expression than the other haplotype, is known, and *M*_*ij*_ is the read count for the corresponding allele that resides on the major haplotype. *M*_*ij*_ is assumed to follow a binominal distribution, *Binomial*(*Y*_*ij*_,*P*_*i*_), where P_*i*_ is the ASE level, representing the underlying transcript frequency of the major haplotype for individual *i* When there is no gene-level ASE, *P*_*i*_ = 0.5, and *P*_*i*_ > 0.5 otherwise. The allele-specific read counts of the major haplotypes are then aligned across all individuals. To account for correlations across multiple *tSNP*s, we employ a generalized linear mixed-effects model in which the random effect is modeled as

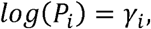

where

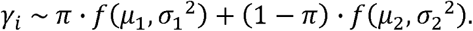

The random effect, *γ*_*i*_, represents the individual-specific true underlying transcript frequency of the distributions with means *μ*_1_ and *μ*_2_, and standard deviations *σ*_1_ and *σ*_2_, respectively. Here, *μ*_1_ and *μ*_2_ major haplotype on a logarithmic scale, whose distribution is assumed to be a mixture of two unknown indicate the population-level major allele transcript frequency of individuals that are heterozygous and homozygous for the unobserved *rSNP* accordingly. Therefore *μ*_1_ represents the underlying true ASE effect which will be deviating from *log* (0.5), whereas *μ*_2_ represents the situation that there is no ASE which will be around *log* (0.5). Parameters are estimated using the non-parametric maximum likelihood estimation (NPML) approach, an Expectation-Maximization based method developed by Murray Aitkin.^16, 17^

To detect gene-level ASE in the population, we test the following hypothesis

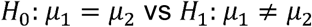

using test statistic 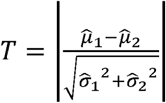. We assess the statistical significance of *T* through a resampling-based procedure. Specifically, for each individual *i* at *tSNP j*, we resample *M*_*ij*_ from *Binomial* (*Y*_*ij*_, 0.5), and calculate the corresponding *T*_*n*_ using the resampled data. We repeat this procedure *N*_*sim*_ times, and gene-specific p-value is calculated as 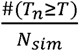.

In the above framework, we have assumed the haplotype phase is known and the major allele can be inferred. However, in real studies, the haplotype phase is often unknown and the observed data offer little or no information of which allele is the major allele. In the absence of DNA genotype data, with only the *X*_*ij*_ and *Y*_*ij*_ of the *tSNP*, it is challenging to infer which alleles at different SNPs reside on the same haplotype. Even when haplotype phase is known, lacking information of the *rSNP*, makes it difficult to align read counts across individuals as we do not know which allele resides on the same haplotype with the *R* allele. To overcome the above-mentioned challenges and determine the major allele, we adapted a pseudo phasing procedure, originally employed by MBASED.^14^ This procedure uses a ‘majority voting’ approach based on observed read counts. For each individual, when the haplotype phase information is known, we assign the haplotype with larger total reads, obtained by summing up read counts across all SNPs on the same haplotype, as the major haplotype. When haplotype phase is unknown, we assign the allele with larger read counts of each SNP to the major haplotype, and alleles on the inferred major haplotype are treated as major alleles.

### Detection of differential ASE between two conditions

The previously described ASE detection procedure for one condition can be naturally extended to detect gene-level ASE difference between two conditions (e.g., conditions A and B) using paired RNA-seq data, where the same individual is sequenced under both conditions. Similar to the one condition analysis, for individual *i* at *tSNP j*, let 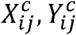 and 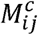 be the condition-specific reference allele read count, total read depth, and major allele read count accordingly for condition 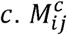 is assumed to follow 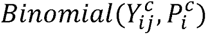, where 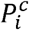 is the condition-specific true underlying transcript frequency of the major haplotype for individual *i* After aligning major alleles across individuals, by introducing a covariate of condition indicator 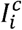, defined as

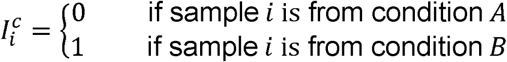

the model can be modified as the following:

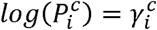

where

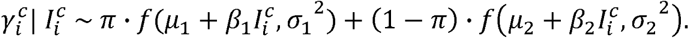

Here 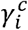 represents 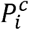 on the logarithmic scale whose conditional distribution is assumed to be a mixture of two unknown distributions, with *μ*_1_ and *μ*_2_ representing the population-level transcript frequency of the major haplotype under condition *A, β*_1_ and *β*_2_ representing the difference in the transcript frequency between the two conditions, for ‘Hom’ and ‘Het’ individuals, respectively. Similarly, parameters are estimated using the NPML approach.^16, 17^ To test gene-level ASE difference between two conditions, we consider the following hypothesis:

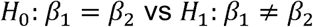

Similar to the one condition analysis, the haplotype phase and major haplotype information are often unknown in real studies. Again, we employ the pseudo phasing procedure to differentiate the major haplotype and align them across individuals.^14^ To ensure that major haplotypes are identical for the same individual under different conditions, we choose one condition as ‘reference’, obtain its phasing information, and phase the data from the other condition accordingly. Ideally, to improve the phasing accuracy, the condition with larger ASE effect is preferred to be used as the ‘reference’.

We consider 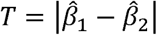 as the test statistic and assess its significance through resampling. To obtain the null distribution of *T*, for each individual *i* at *tSNP j*, we resample 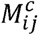 from 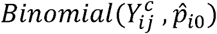, where 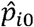 is the individual-specific estimate of the ASE level assuming no ASE difference between the two conditions. A two-step procedure is employed to obtain 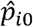. First, for individual *i*, we combine data from both conditions, and calculate 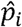 as the transcript frequency of the major haplotype in the pooled sample, where

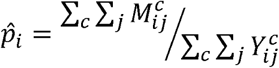

Because we perform pseudo alignment on the RNA-seq data based on a ‘reference’ condition, after the ‘majority voting’, for ‘Hom’ individuals, 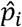, as the pooled major allele frequency, will always be larger than 0.5, which violates our assumption of no ASE effect under both conditions. To make the resampled data represent the null, as a second step, 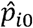 is obtained through a weighted sum as the following:

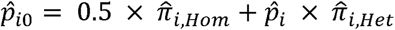

where 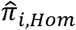 and 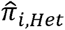 are the estimated posterior probabilities that individual *i* belongs to the ‘Hom’ and ‘Het’ group, respectively. Based on the resampled data, *T*_*n*_ can be obtained accordingly. This procedure is repeated *N*_*sim*_ times, and the p-value is calculated as 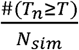.

### Simulation framework

Without loss of generality, we consider one gene only. To evaluate the performance of ASEP across a wide range of scenarios, we simulated RNA-seq data for *N* individuals (20 or 50), each with *nSNP* the number of *tSNP*s (2, 4 or 6). For each individual, we generated the data with a pre-specified minor allele frequency (*MAF*) of the rSNP (0.1, 0.3 or 0.5), and assigned ‘Hom’ or ‘Het’ based on the genotype of the *rSNP*. The haplo-genotype data were simulated assuming Hardy-Weinberg equilibrium (HWE) with assigned haplotype frequencies so that, for each tSNP, *MAF* = 0.3 with the LD coefficient between pairs of *tSNP*s set at 0.8.

### Simulation scheme for ASE detection under one condition

The read count for the major allele of each *tSNP* was sampled from *Binomial* (*Y*_*ij*_, 0.5) for ‘Hom’ individuals and from *Binomial* (*Y*_*ij*_, *P*_*i*_) for ‘Het’ individuals across all simulations. For simplicity, we assume *Y*_*ij*_ = *Y* for all individuals across all *tSNP*s, where *Y* takes two possible values, 50 or 100. For ‘Het’ individuals, when evaluating the type ? error rate, we set *P*_*i*_ = 0.5 under both phase known and unknown scenarios. When evaluating power, to account for subject-specific random variation in ASE levels, *P*_*i*_ was simulated from *N* (*P*,0.03^2^), where *P* is the pre-specified ASE effect in the population. We set *P* = 0.55 under phase known situation and *P* = 0.6, otherwise.

### Simulation scheme for differential ASE detection between two conditions

Similar to one condition analysis, for ‘Hom’ individuals, the major allele read count for each *tSNP* was simulated from 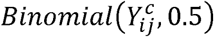 for both conditions across all evaluations. For ‘Het’ individuals, the major allele read count was simulated from 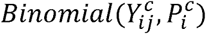, where *c* represents condition (*A* or *B*). For simplicity, we assume 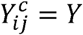, where *Y* takes two values 100 or 200. When evaluating the type I error rate, we set 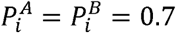 under both phase known and unknown scenarios. When evaluating the power, 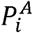 and 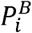 were sampled from *N* (*P*^*A*^,0.03^2^) and *N* (*P*^*B*^,0.03^2^), respectively, where *P*^*A*^ and *P*^*B*^ are the pre-specified condition-specific ASE effect in the population for condition *A* and condition *B*. When haplotype phase is known, we set 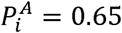 and 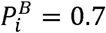, and 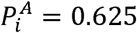 and 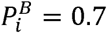, otherwise. Condition *B* was used as the ‘reference’ for pseudo-phasing given its stronger ASE effect.

### Human macrophage differentiation and polarization

All of the protocols for this study were approved by the Human Subjects Research Institutional Review Board at the University of Pennsylvania and Columbia University Irving Medical Center. Peripheral blood mononuclear cell (PBMC) collected using BD VACUTAINER® Mononuclear Cell Preparation Tube were cultured in macrophage culture media, 20% FBS in RPMI 1640 media with 100 ng/ml human macrophage colony-stimulating factor (M-CSF), for 7 days on BD Primaria™ tissue culture plate to induce macrophage differentiation.(PMID: 25904599^18^, 28882870^19^) Polarization was obtained in the presence of M-CSF by 18-20 hour incubation with 20 ng/ml interferon-gamma (IFN-*γ*) and 100 ng/ml lipopolysaccharide (LPS) for M1-like polarization. (PMID: 25904599^18^, 28882870^19^)

### RNA-Seq library preparation and sequencing

RNA samples of M0 and M1 macrophages were extracted using All Prep DNA/RNA/miRNA Universal Kit (Qiagen, Valencia, CA) (PMID: 25904599^18^, 28882870^19^) by batches and the samples were randomly assigned to each batch. The RNA quality and quantity were determined by Agilent 2100 Bioanalyzer (Median RIN = 7.9, n = 96 samples from 48 subjects). With a minimum of 300 ng input RNA, libraries were prepared using the TruSeq RNA Sample Preparation Kit (Illumina, San Diego, CA), followed by 101 bp 60M paired-end sequencing on an Illumina’s HiSeq 4000 at Columbia Genome Center.

## RESULTS

### Detecting ASE under one condition

We evaluated the performance of ASEP to detect gene-level ASE under one condition across a wide range of scenarios. We first considered the situation when phase among the *tSNPs* is known. Our simulations showed that type I error rate was controlled at the 5% level under all scenarios we investigated. As expected, the power of ASEP increases as the number of individuals, sequencing depth, or the number of heterozygous *tSNPs* increases. Among these three factors, the sequencing depth is the most influential one as compared to the other two. In addition, the magnitude of power increase for a specific factor depends on the values of the other factors. For example, the effect of sequencing depth or sample size is minimal until the number of *tSNP*s increases from 2 to 4, and the effect becomes much stronger when the number of *tSNPs* increases further to 6. With large sample size, high sequencing depth and more *tSNP*s, our method has sufficient power to detect ASE effect that is as small as 0.05. Further, increasing the proportion of ‘Het’ individuals in the sample, determined by the *MAF* of the *rSNP*, improved the model performance under most of the scenarios considered. The model performed similar with *MAF* = 0.3 or *MAF* = 0.5, and outperformed the same model with *MAF* = 0.1. This is expected since when *MAF* = 0.1, only 18% of individuals are heterozygous at the *rSNP* under HWE, whereas more than 40% of the individuals are heterozygous with *MAF* = 0.3 or 0.5, leading to larger effective sample size in detecting ASE (**Figure 2A** and **Figure S1A**).

**Figure 2.**
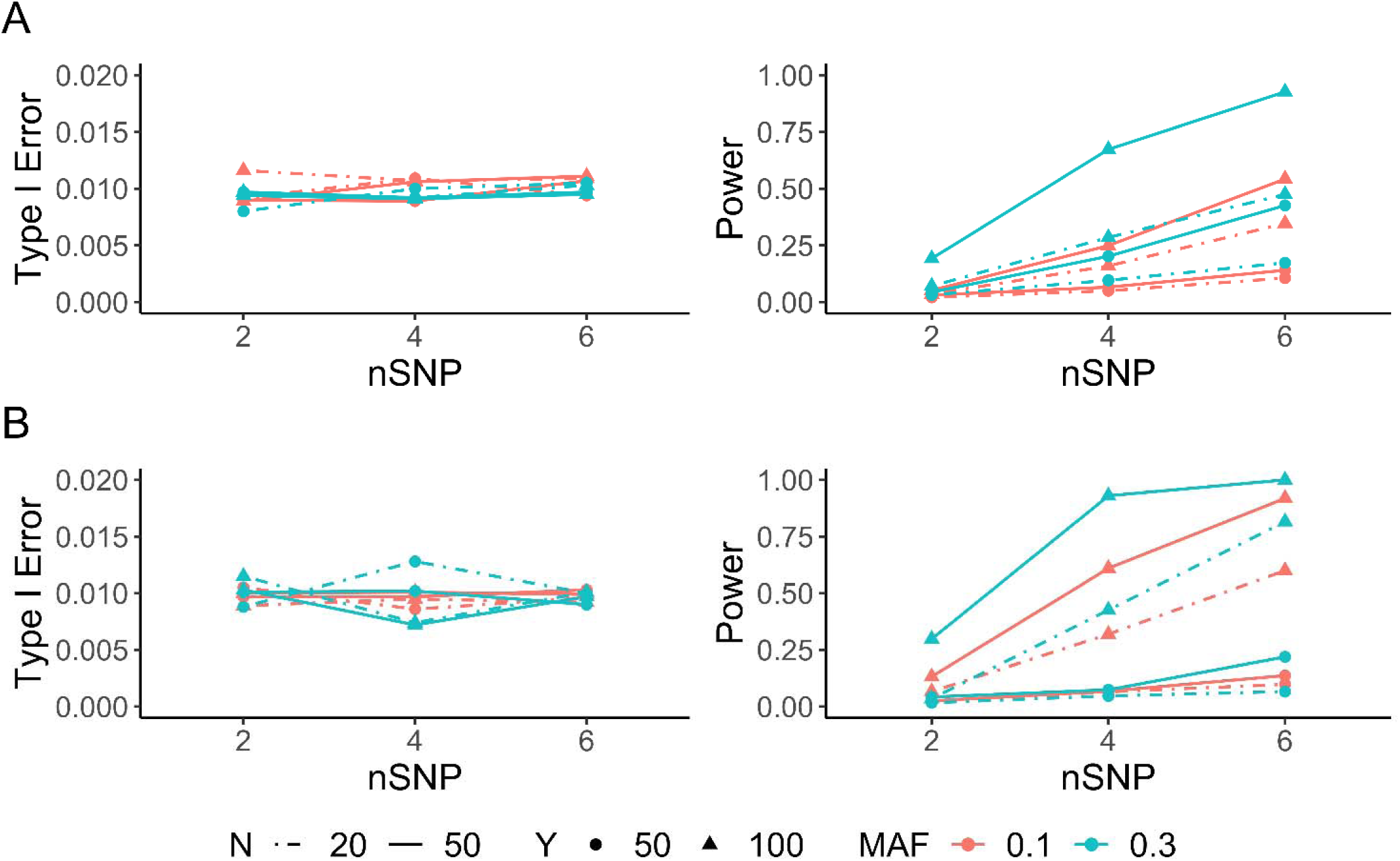
Simulation results for one-condition analysis. Type I error rate (left) and power (right) evaluated as a function of the number of individuals (*N*), sequencing depth (*Y*), number of heterozygous transcribed SNPs (*nSNP*) and *MAF* of the *cis*-regulatory SNP. For each scenario, the type I error rate was examined with 10,000 simulations, and the power with 1,000 simulations at significance level *α* = 0.01. Performance of ASEP **(A)** when haplotype phase is known and the population-level ASE is 0.55 for power evaluation and **(B)** when haplotype phase is unknown and the population-level ASE is 0.6 for power evaluation.

Next, we examined the performance of ASEP when haplotype phase is unknown. The type I error rate was still under control across all scenarios considered. For power evaluation, due to potential phasing errors caused by separating paternal and maternal alleles, we increased the true underlying ASE effect from 0.55 to 0.6 in order to have adequate detection power. As expected, the read depth is still the most important factor as compared to sample size and the number of *tSNP*s. Additionally, increasing the proportion of ‘Het’ individuals leads to improved power under all scenarios, similar to what we observed for phase known situation (**Figure 2B** and **Figure S1B**).

### Detecting differential ASE between two conditions

We then evaluated the performance of ASEP to detect ASE difference between two conditions. When haplotype phase information is known, the type I error rate of ASEP was well controlled across a variety of settings. Similar to the one condition analysis, when *MAF* of the *rSNP* is fixed, the power increased as the number of individuals, sequencing depth, or the number of *tSNP*s increased. Among these three factors, read depth had the largest impact on power. Moreover, a specific factor only had minor impact on power when the other two factors were at low level. When any two of the three factors were at high level, ASEP had adequate power to detect a 0.05 ASE difference between two conditions. However, unlike one condition analysis, increasing *MAF* of the *rSNP* did not necessarily improve the model performance for detecting differential ASE between two conditions. In fact, its effect depended on the value of the three aforementioned factors. Power increased with *MAF* only when number of *tSNP*s are high or number of *tSNP*s are moderate but with high sequencing depth (**Figure 3A** and **Figure S2A**).

**Figure 3.**
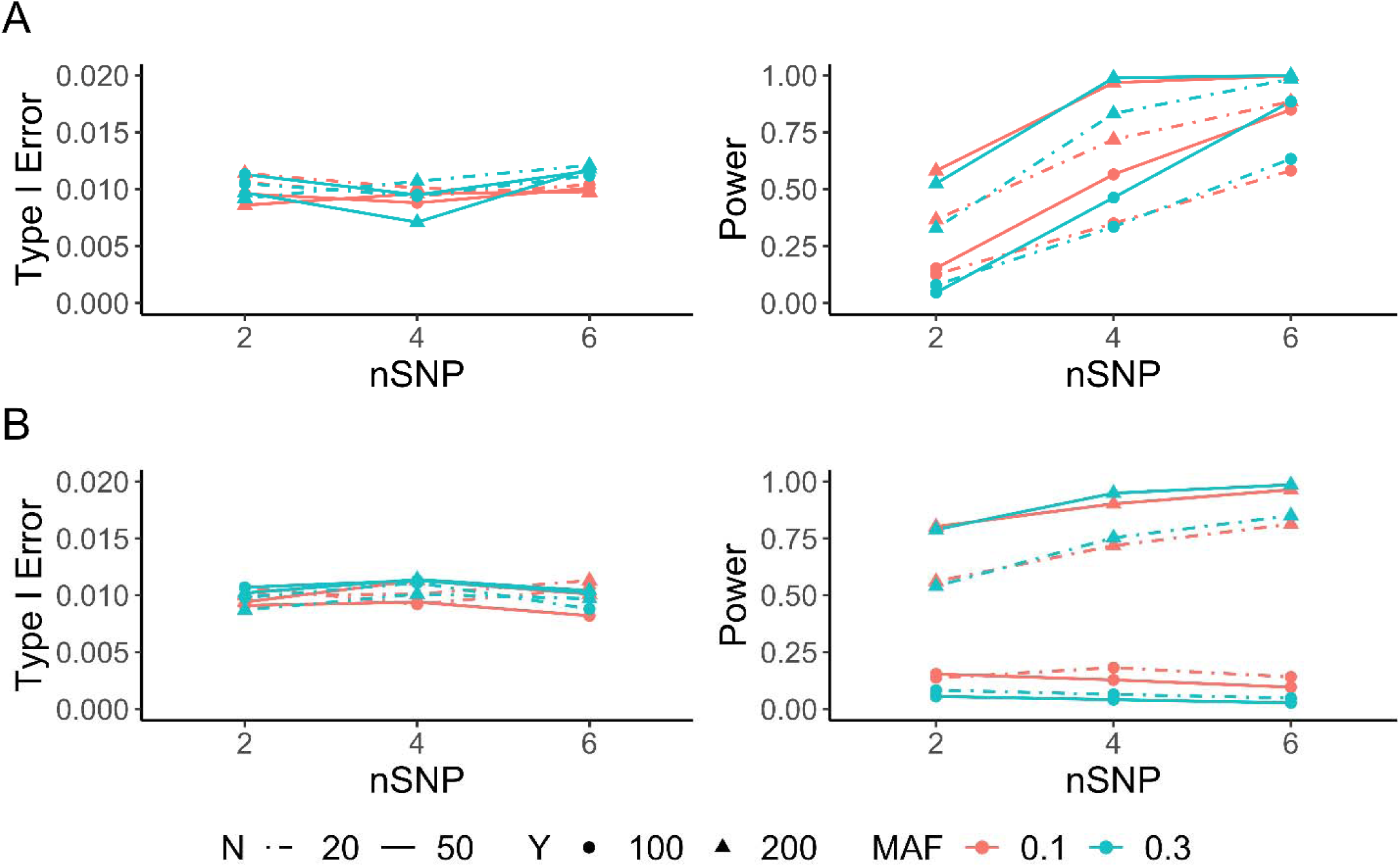
Simulation results for two-condition analysis. Type I error rate (left) and power (right) evaluated as a function of the number of individuals (*N*), sequencing depth (*Y*), number of heterozygous transcribed SNPs (*nSNP*) and *MAF* of the *cis*-regulatory SNP. For each scenario, the type I error rate was examined with 10,000 simulations, and the power with 1,000 simulations at significance level *α* = 0.01. **(A)** Performance of ASEP when haplotype phase is known. For power evaluation, the population-level ASE takes values of 0.7 and 0.65 for the two conditions. **(B)** Performance of ASEP when haplotype phase is unknown. For power evaluation, the population-level ASE takes value of 0.7 and 0.625 for the two conditions.

When the haplotype phase is unknown, the type I error rate was still under control across all scenarios. For power evaluation, as compared to phase known situation, we increased the ASE difference between two conditions from 0.05 to 0.075 in order to have adequate power. At a fixed level of *MAF*, the power only increased as sequencing depth increased. For the other two factors, the number of individuals and number of *tSNPs*, they improved the power only when sequencing depth was high. This is not surprising since when phase is unknown, the phasing errors increase as the number of individuals or the number of *tSNPs* increases. Interestingly, *MAF* overall had a negative effect on the model performance under most of the situations. It only improved the model performance with the combination of high read depth, large sample size, and moderate to high number of SNPs (**Figure 3B** and **Figure S2B**).

### Application to a human macrophage RNA-seq dataset

We applied ASEP to a paired macrophage RNA-seq dataset generated from 48 healthy individuals (Supplementary Table S5 for subject demographics). Human PBMC can be cultured and differentiated to macrophages, and polarized *in vitro* to functionally and molecularly distinct M1-like inflammatory macrophages by IFN-γ and LPS, an important and widely-used experimental model to study macrophage biology in homeostasis and diseases (PMID: 26982353^20^, 30679807^21^). M0 and M1 macrophages from each individual were subject to 2×101 bp paired-end RNA-seq. Reads were aligned to human genome hg19 using STAR 2.6.0a^22^. Reads from each pair were required to map to the same chromosome with distance <500,000 bp. Only uniquely mapped reads were retained for downstream analysis. The RNA-seq data were processed using WASP^23^ to remove the possible mapping bias and extract allele-specific read counts.

We first applied ASEP for one condition analysis to M0 and M1 macrophage samples separately to detect ASE genes under each condition. For a given gene, a *tSNP* is included in the analysis if the minor allele count ≥ 5, total read depth ≥ 20, and the minor allele count is at least 5% of the total read count. Genes that appeared in less than three individuals were excluded from analysis. In total, we analyzed 5,961 genes for the M0 samples and 5,465 genes for the M1 samples, with 4,783 genes common in both samples. We identified 792 genes with significant ASE (*P* < 0.05) in M0 and 747 genes in M1. Among these genes, 634 were detected only in M0, with 124 of them not expressed in M1, and 589 genes detected only in M1, with 102 of them not expressed in M0. Additionally, 158 genes were found to show ASE under both conditions. After multiple testing adjustment with false discovery rate (FDR), 160 genes remained significant (adjusted *P* < 0.05) in M0 and 219 genes in M1 (**Tables S1** and **S2**). Additionally, 141 genes were detected only in M0 with 16 of them not expressed in M1, 200 genes were detected only in M1 with 28 of them not expressed in M0, and 19 genes were found to have ASE under both conditions (**Figure 4A**). For those significant genes, we further performed gene set enrichment analysis using metascape.^24^ ASE genes detected in M0 are enriched for genes related to adaptive immune system and autophagy, whereas those detected in M1 are enriched for genes related to cytokine signaling in immune system and cellular responses to external stimuli (**Figure 5A**).

**Figure 4.**
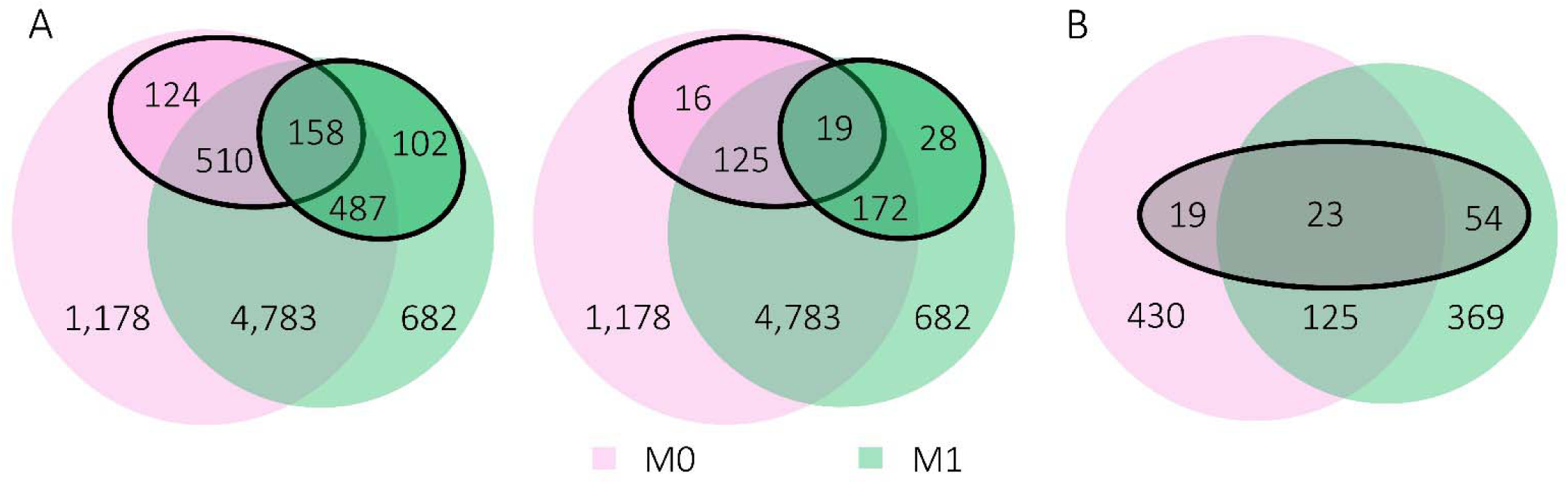
Genes analyzed for ASE and differential ASE analysis in the macrophage RNA-seq dataset. **(A)** Total number of genes analyzed, and number of significant ASE genes in M0 (pink) and M1 (green) macrophages obtained from one-condition analysis. Solid circles indicate the nominal significant (left) and FDR-adjusted significant (right) ASE genes detected under each condition. **(B)** Total number of genes analyzed for two-condition analysis. Genes were selected from significant (nominal) ASE genes for M0 (pink) and M1 (green) macrophages. Solid circle indicates the FDR-adjusted significant differential ASE genes detected between M0 and M1.

**Figure 5.**
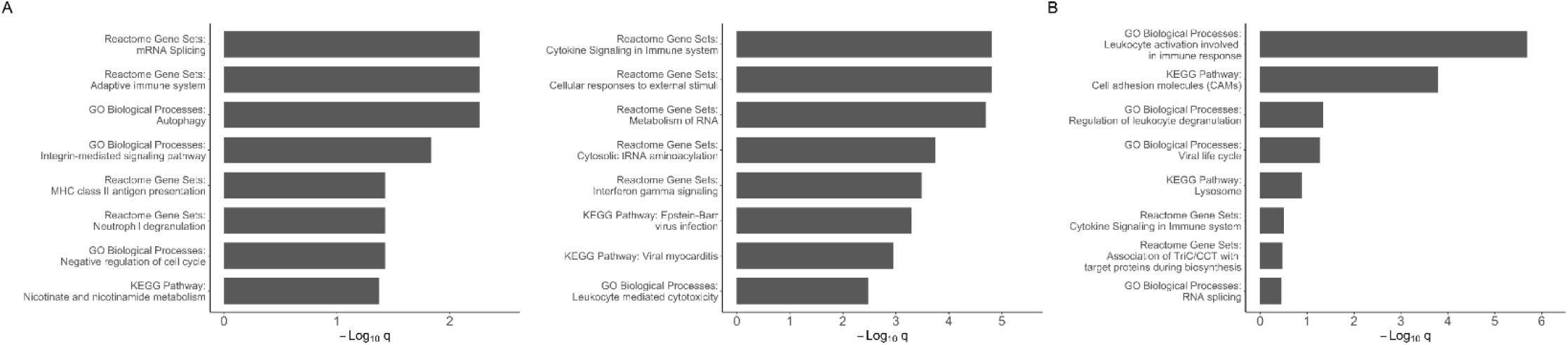
Representative enriched terms of ASE and differential ASE genes in the macrophage RNA-seq dataset. **(A)** Top representative enriched terms for significant ASE genes for M0 (left) and M1 (right) samples. **(B)** Top representative enriched terms for significant differential ASE genes between M0 and M1 samples. The q-values were calculated using Banjamini-Hochberg multiple testing adjustment.

Encouraged by the above analysis results, we next performed the differential ASE analysis between M0 and M1 by selecting 924 candidate genes from one condition analysis that were found to show evidence of ASE (*P* < 0.05) in M0 or M1. Since haplotype phase is unknown, to reduce phasing error, for each gene, we chose the condition with higher estimated ASE effect as the ‘reference’ to phase the data from the other condition. In total, we detected 190 genes showing evidence of differential ASE (*P* < 0.05), with 96 genes still being significant after multiple testing adjustment (FDR adjusted *P* < 0.05) (**Figure 4B** and **Table S3**). We performed enrichment analysis for these 96 genes, and as expected, these differential ASE genes are highly enriched for leukocyte activation involved in immune response (**Figure 5B**).

Since macrophages are important regulators and promoters of many cardiovascular disease programs, next, we examined whether these 96 genes showing significant differential ASE overlap with findings from GWAS for cardiovascular disease (CVD), coronary artery disease (CAD), and acute coronary syndrome (ACS).^25^ Among the 96 genes, 52 overlapped with loci that reached GWAS significance (*P* < 5 × 10^−8^) (**Table S4**). For example, *PECAM1* showed strong evidence of differential ASE (FDR adjusted *P* < 0.0001). A previous study showed that *PECAM1* is up-regulated by LPS stimulation, and may function as a feedback negative regulator of inflammatory response in macrophages.^26^ **Figure 6** shows the estimated SNP-level ASE difference for each individual, i.e., the difference in major allele proportion of each SNP after haplotype phasing between M1 and M0 samples for each individual. After sorting individuals by their median of estimated ASE difference among heterozygous SNPs, we observed that, majority of the individuals have positive ASE difference with a few having negative ASE difference, which might be due to potential phasing uncertainty. About one-third of the subjects have median ASE difference around zero, which presumably are individuals with homozygous *cis*-regulatory SNP. However, since more individuals have positive ASE difference, our method was able to detect evidence of differential ASE at the population level by aggregating information across individuals and transcribed SNPs (**Figure 6A**).

**Figure 6.**
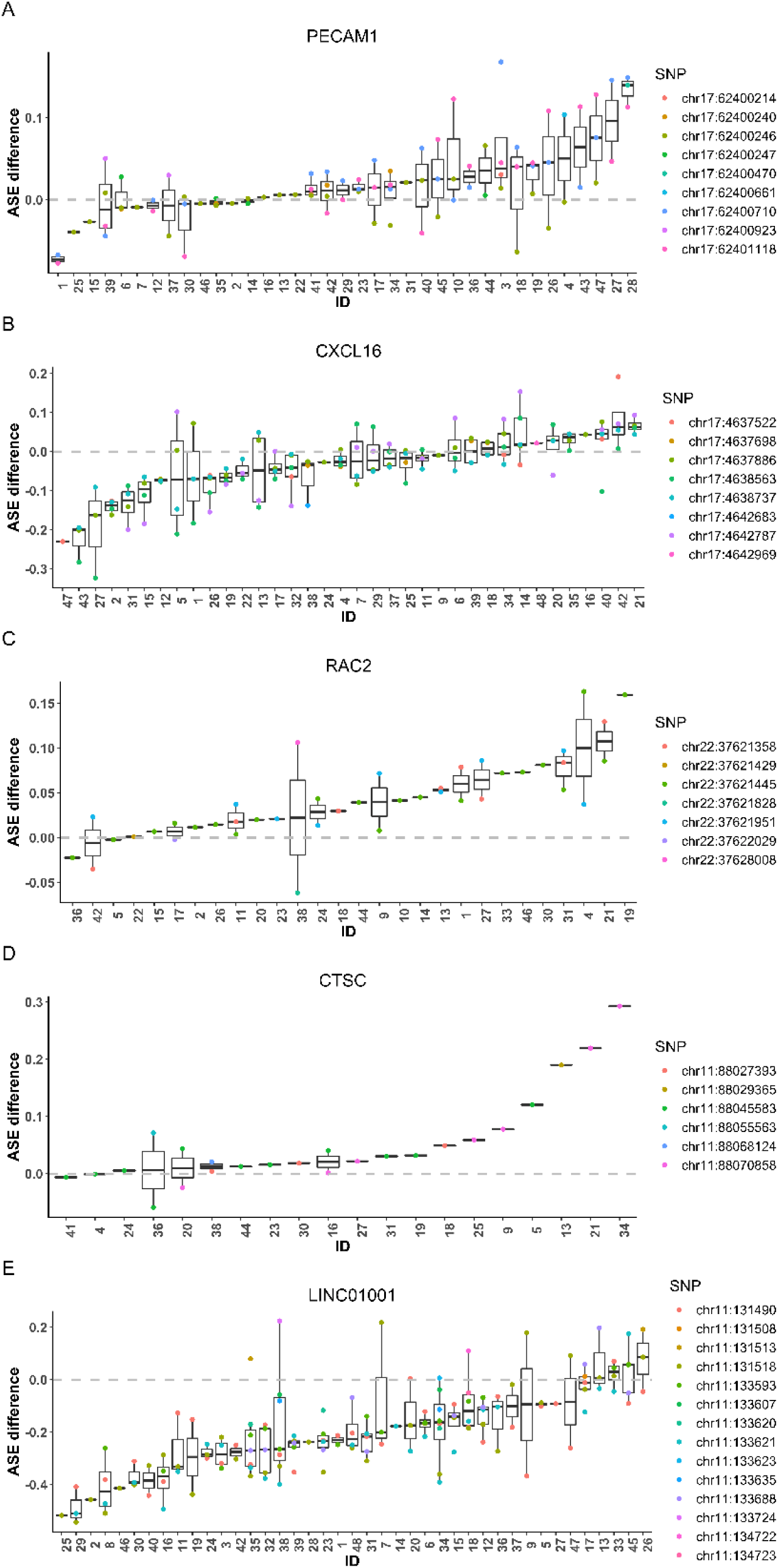
SNP-level ASE difference between M0 and M1 macrophage samples for selected genes showing differential ASE in the macrophage RNA-seq dataset. We selected five genes, *PECAM1* **(A)**, *CXCL16* **(B)**, *RAC2* **(C)**, *CTSC* **(D)** and *LINC01001* **(E)**, to show their estimated SNP-level ASE difference for each SNP and individual. The estimated ASE difference was calculated as the difference in the major allele proportion between M1 and M0 samples after haplotype phase alignment. The individuals were sorted by the median ASE difference across all SNPs of each individual.

We also detected differential ASE in *CXCL16* (FDR adjusted *P* = 0.0007) in the macrophage data, which has been reported to play a critical role on inflammation. *CXCL16* acts as a scavenger receptor and promotes the uptake of modified lipids that leads to foam cell formation.^27^ Lehrke *et al*. showed that *CXCL16* may associate with human atherosclerosis, and is correlated with inflammatory and metabolic response of acute coronary syndrome.^25^ Our RNA-seq data suggest that approximately half of the subjects have estimated ASE difference around zero, whereas the other half have ASE difference deviating considerably from zero. By accounting for sample heterogeneity, ASEP was able to uncover differential ASE between conditions (**Figure 6B**).

Additionally, we detected differential ASE for *RAC2* (FDR adjusted *P* = 0.0026), an essential element in regulating B lymphocyte functions and the corresponding immune response.^28^ Although each individual only has a few SNPs, by accumulating evidence across multiple individuals, we were able to detect a consistent signal of differential ASE (**Figure 6C**). Further, we detected differential ASE in *CTSC* (FDR adjusted *P* = 0.0039), which encodes cathepsin C, a central activator of many serine proteases in immune and inflammatory cells.^29^ Heterozygous sites in *CTSC* varied across individuals and most of them are heterozygous for only one SNP in this gene. Although individuals are heterozygous for different SNPs, for each SNP, the ASE effect was consistently larger for the M1 samples. Therefore, by aggregating information across multiple SNPs and individuals, ASEP was able to detect a population-level differential ASE effect (**Figure 6D**).

Other genes that showed significant differential ASE include those from the human leukocyte antigen (HLA) gene complex (*HLA-DQA2, HLA-F* and *HLA-B* (FDR adjusted *P* < 0.0001)),^30^ *CD4* (FDR adjusted *P* = 0.0056), *SIGLEC1* (FDR adjusted *P* = 0.0093), *HMGCS1* (FDR adjusted *P* = 0.015), *ACAP2* (FDR adjusted *P* = 0.0162) (**Figure S3**).

Although 44 of the differential ASE genes did not overlap with GWAS findings, they also seemed to be interesting and may play a role relevant in inflammation. For example, *LINC01001*, a long intergenic noncoding RNA (lincRNA), showed strong evidence of differential ASE (FDR adjusted *P* < 0.0001) (**Figure 6E**), and it might be involved in coordinated response during macrophage activation, and potentially regulates the inflammatory response in cardiometabolic disease.^31^ Other genes of interest include *NOTCH2* (FDR adjusted *P* < 0.0001), *LILRB3* (FDR adjusted *P =* 0.0006), and *TLN1* (FDR adjusted *P =* 0.0056) (**Figure S3**). *NOTCH2* plays a role in notch signaling that regulates macrophage activation in pro-inflammatory process.^32^ *LILRB3* encodes a protein that belongs to the leukocyte immunoglobulin-like receptor (LIR) family, where it binds to MHC class I molecules on antigen-presenting cells and acts as an inhibitory receptor of the immune response.^33^ *TLN1* encodes a cytoskeletal protein, talin-1, that has been implicated in phagocytosis, as well as regulates T cell proliferation and functions.^34, 35^

## DISCUSSION

Detecting ASE is an important step towards the understanding of genetic polymorphisms on gene expression variation. However, existing ASE detection methods mainly focused on individual-based ASE effect. To better utilize shared ASE information across individuals, we proposed ASEP, a novel method that can detect allelic imbalance in gene expression across individuals under one condition, and differential ASE between two conditions. The main advantage of ASEP lies in its ability to leverage information across multiple individuals and SNPs within the same gene. Existing methods, such as MBASED^14^, detect ASE effect through individual-based analysis, which makes it difficult to aggregate information across subjects. GeneiASE^15^ uses Fisher’s meta-analysis method to combine p-values across subjects, however, the resulted p-value is driven by extremely small p-values, which may lead to a significant combined p-value even when ASE is absent in the majority of the subjects.

A major challenge for cross-individual ASE analysis is due to the difficulty in differentiating ‘Hom’ and ‘Het’ individuals as the underlying *cis*-rSNP is unobserved. By employing a mixture model, ASEP is able to aggregate ASE effects contributed by those ‘Het’ individuals while accounting for the heterogeneity introduced by those ‘Hom’ individuals. As a result, ASEP is not only more powerful but its result is also easier to interpret compared to traditional individual-based ASE tests. Through extensive simulations, we showed that ASEP is sensitive in detecting small ASE effect under a wide range of scenarios. We further demonstrated that the ASE effects uncovered by ASEP are convincing through the analysis of a RNA-seq dataset on human macrophages. The detected ASE and differential ASE genes between M0 and M1 samples agreed well with relevant biological pathways and previous GWAS findings.

ASEP can be applied when haplotype phase information is known or unknown. When sequencing depth is high, the haplotype phase reconstruction approach employed by ASEP is able to correctly recover the true major haplotype. For genes with relatively low sequencing depth, correct assignment of haplotype phase will increase the power to detect ASE. Since *rSNP* is unobserved, paired RNA-seq data are needed for two-condition analysis in order to correctly phase the haplotypes and aligned them consistently across samples from both conditions. If genotype data and phase information are available, then based on the alleles of the regulatory SNP which is either known or hypothetical, we not only can differentiate the ‘Het’ individuals but also can easily align ‘major’ haplotypes which resides on the same haplotype with the expression-increasing allele across individuals. This way, our method can be further modified to detect ASE difference using all available data or even for unpaired samples, such as case-control study, to detect the differential ASE between two independent groups. ASEP is a regression-based framework for ASE analysis, which is flexible and can be easily extended to adjust for additional covariates or confounders in the model if necessary. Second, as a method designed for analysis of bulk RNA-seq data, we cannot tell if the detected ASE is driven by cell type composition change or cell type-specific ASE. Therefore, for future study, investigating cell-type specific ASE will help provide extra information and will be more powerful especially for genes expressed in rare cell types.

In summary, we have developed ASEP, a gene-based ASE detection method by aggregating information across individuals and SNPs within the same gene. ASEP can detect genes with shared ASE effect or differential ASE across individuals in a population, which leads to easier interpretation and improved power as compared to traditional individual-based ASE detection methods. With the wide application of RNA-seq in biomedical studies, more samples of the same tissue from different individuals are being sequenced in order to study gene-phenotype correlation. There is an urgent need to learn a comprehensive picture of ASE in the broad population instead of focusing on individual-level effect. We believe ASEP, which, to the best of our knowledge, is the first method for population-based ASE detection, will be well-suited for various ASE studies for human diseases.

## ACKNOWLEDGEMENTS

This work was supported by the following grants: NIH R01GM108600 and R01GM125301 (to M.L.), R01HL113147 (to M.L. and M.P.R.), R00HL130574 (to H.Z.), and pilot grant through UL1TR001873 (to H.Z.).

## WEB SOURCES

ASEP R package is available on Github (https://github.com/Jiaxin-Fan/ASEP).

