## Supplemental Data for "ASEP: gene-based detection of allele-specific expression in a population by RNA-seq"

**Supplementary Methods**

**Supplementary Figures**

Figure S1. Simulation results for one condition analysis.

Figure S2. Simulation results for two condition analysis.

Figure S3. SNP-level ASE difference between M0 and M1 macrophage samples for selected genes showing differential ASE in the macrophage RNA-seq dataset.

**Supplementary Tables**

Table S1. Significant ASE genes in M0 macrophage samples.

Table S2. Significant ASE genes in M1 macrophage samples.

Table S3. Significant differential ASE genes between M0 and M1 macrophage samples.

Table S4. Significant differential ASE genes between M0 and M1 macrophage samples that overlap with GWAS loci.

Table S5. Subject demographics of the macrophage samples.

**Supplementary Methods**

**Human macrophage differentiation and polarization** Peripheral blood mononuclear cells (PBMCs) were collected using BD VACUTAINER® CPT™ Cell Preparation Tubes with Sodium Citrate. The freshly isolated PBMC were plated (6-well plates, 3x10^6^ cells per well, or 24-well plates, 1x10^6^ cells per well) (BD Primaria^TM^ cell culture plates, BD Biosciences, Bedford, MA), allowed to adhere (overnight), washed with RPMI-1640 and cultured in macrophage culture media (20% FBS in RPMI 1640 media with 100 ng/ml human M-CSF) (300-25, Peprotech, Rocky Hill, NJ) for 7 days to induce macrophage differentiation as we described. (PMID: 25904599^1^, 28882870^2^). Macrophage polarization was obtained by removing culture medium at Day 6 and culturing cells for an additional 18-20 hour in 10% FBS in RPMI 1640 media with 50 ng/ml M-CSF for M0 control as the unstimulated macrophages, or the above media supplemented with 20 ng/ml IFN-γ (R&D, Minneapolis, MN) and 100 ng/ml lipopolysaccharide (LPS; Sigma-Aldrich, St. Louis, MO) for M1 polarization.

**RNA-seq library preparation and sequencing** As we described (PMID: 25904599^1^, 28882870^2^), RNA samples were extracted using All Prep DNA/RNA/miRNA Universal Kit (Qiagen, Valencia, CA) by batches and the samples were randomly assigned to each batch. The RNA quality and quantity were determined by Agilent 2100 Bioanalyzer using RNA 6000 Nano Kits. With a minimum of 300 ng input RNA, libraries were prepared using the TruSeq Stranded mRNA Library Prep Kit (RS-122-2101, Illumina, San Diego, CA) according to the manufacturer’s protocol with the following modification: 1) the fragmentation time was decreased from 8 to 6 min to ensure libraries were > 100bp long and 2) PCR amplification was limited to 12 cycles for library enrichment to avoid bias from PCR “jackpot” mutations.(PMID: 25904599^1^, 28882870^2^) Library length and concentration were evaluated with the Agilent 2100 Bioanalyzer and PCR quantification (KAPA) and pooled at 2 nM for massively parallel sequencing (2 x 100 bp for 30 million paired-end reads) performed on an Illumina’s HiSeq 4000. (PMID: 25904599^1^, 28882870^2^) On average, we obtained ~101 million filtered reads per sample with ~88% mapping rate.

**Supplementary Figures**

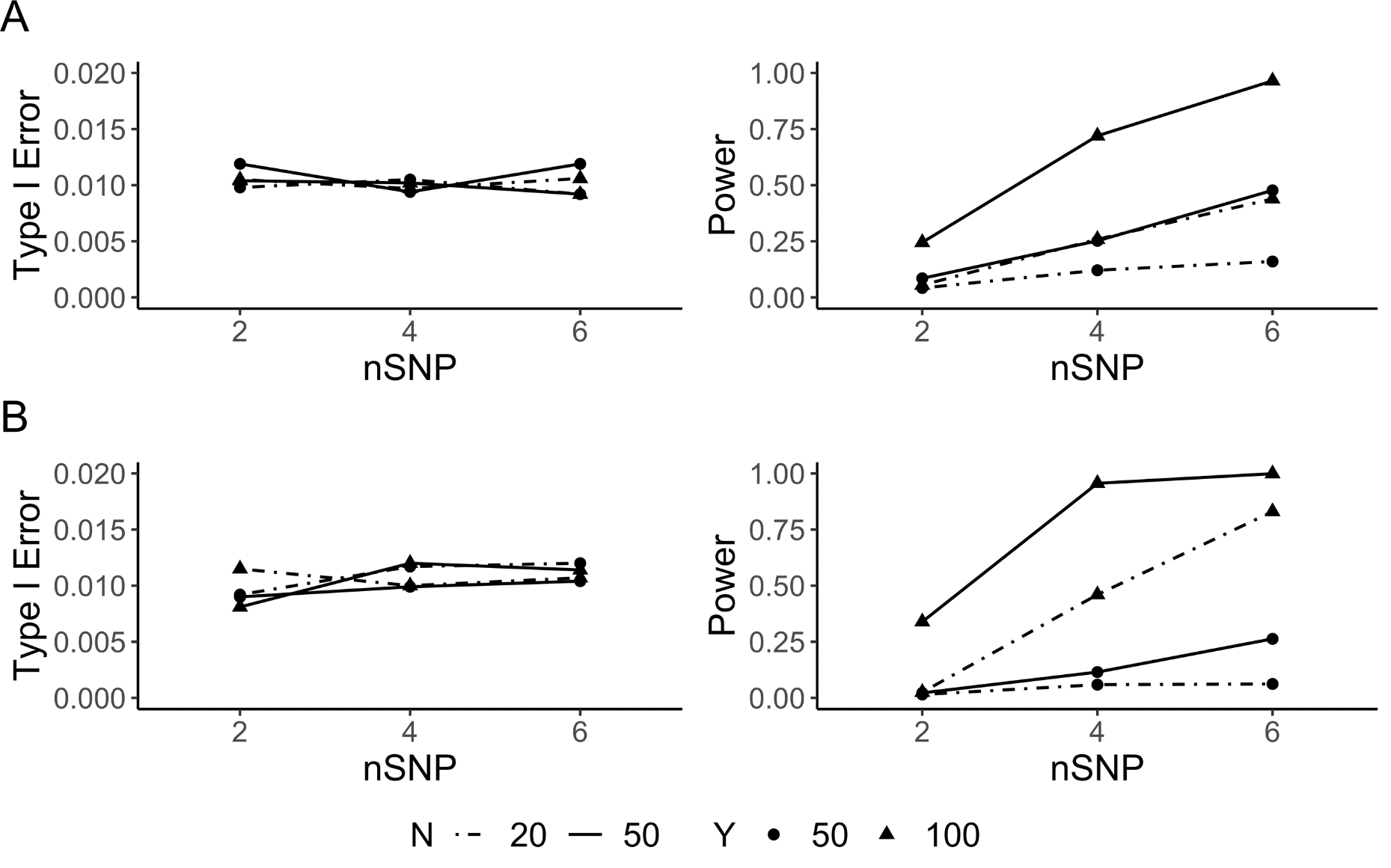

**Figure S1. Simulation results for one-condition analysis**. Type I error rate (left) and power (right) evaluated as a function of the number of individuals ($N$), sequencing depth ($Y$), and the number of heterozygous transcribed SNPs ($nSNP$) when the $MAF$ of *cis*-regulating SNP is 0.5. For each scenario, the type I error rate was estimated based on 10,000 simulations, and the power was estimated based on 1,000 simulations at significance level $\alpha=0.01$. **(A)** Performance of ASEP when haplotype phase is known. The population-level ASE was pre-specified as 0.55 for power evaluation. **(B)** Performance of ASEP when haplotype phase is unknown. The population-level ASE was pre-specified as 0.6 for power evaluation.

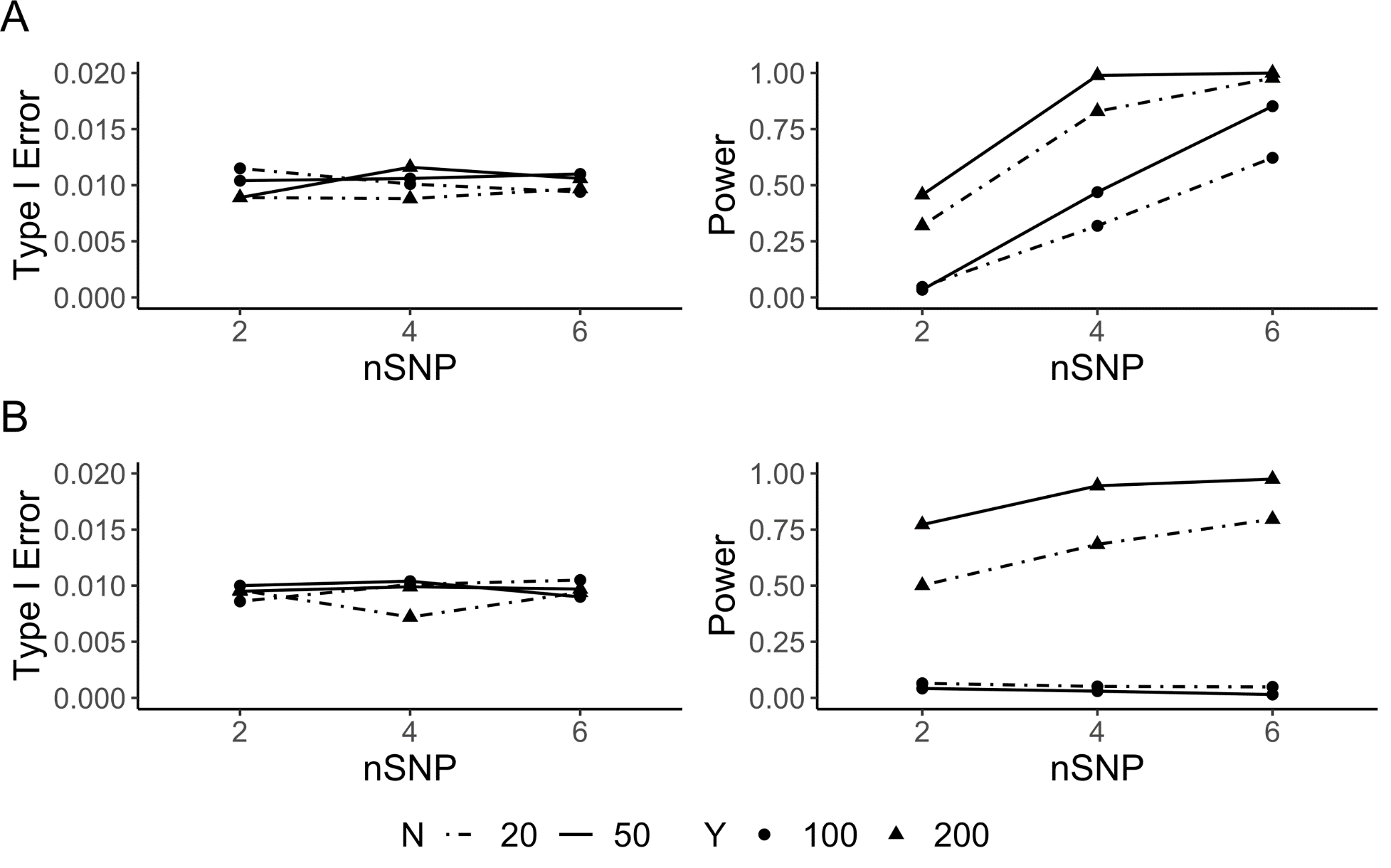

**Figure S2. Simulation results for two-condition analysis.** Type I error rate (left) and power (right) evaluated as a function of the number of individuals ($N$), sequencing depth ($Y$), and the number of heterozygous transcribed SNPs ($nSNP$) when the $MAF$ of *cis*-regulating SNP is 0.5. For each scenario, the type I error rate was estimated based on 10,000 simulations, and the power was estimated based on 1,000 simulations at significance level $\alpha=0.01$. **(A)** Performance of ASEP when haplotype phase is known. For power evaluation, the population-level ASE takes values of 0.7 and 0.65, respectively, for the two conditions. **(B)** Performance of ASEP when haplotype phase is unknown. For power evaluation, the population-level ASE takes values of 0.7 and 0.625, respectively, for the two conditions.

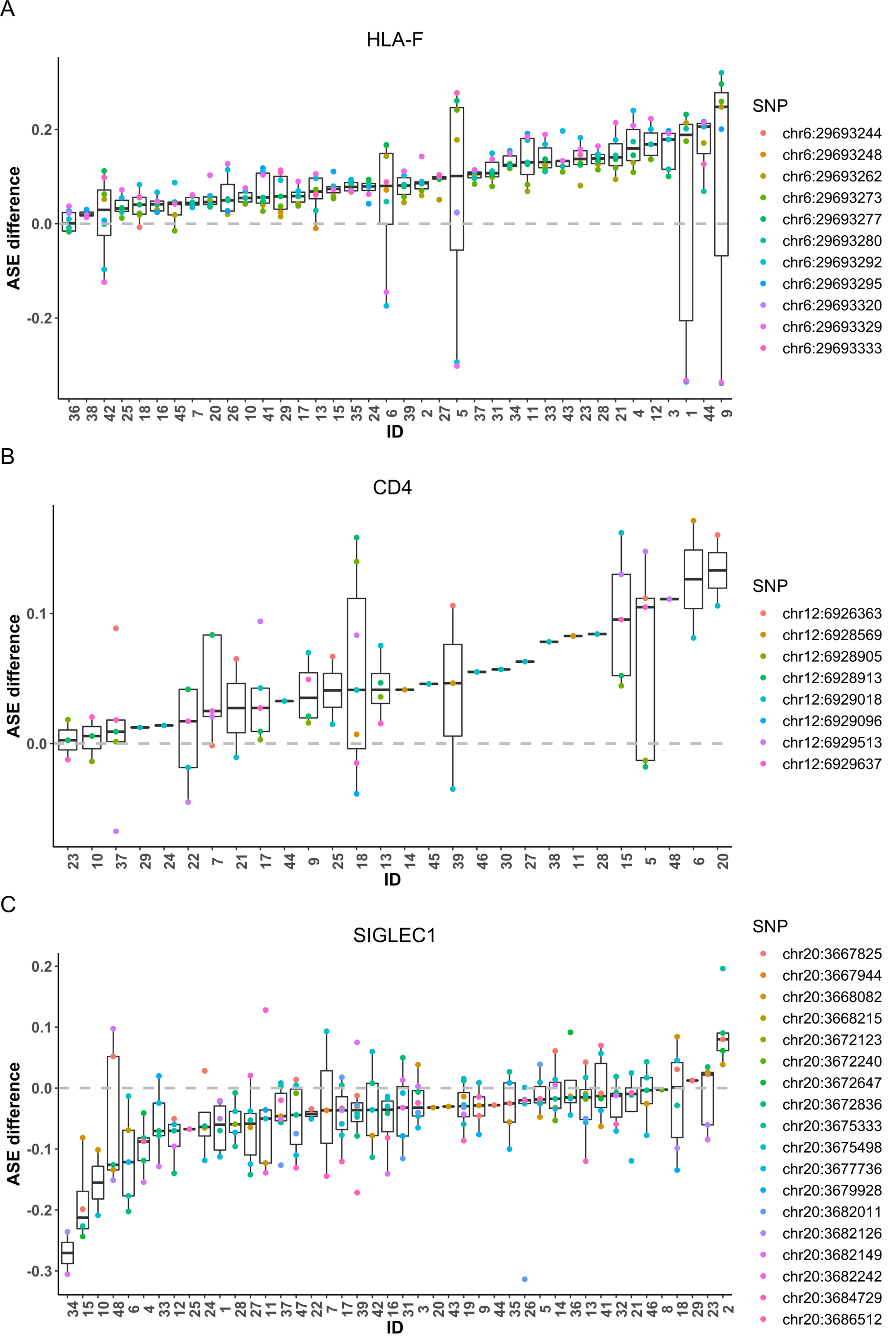

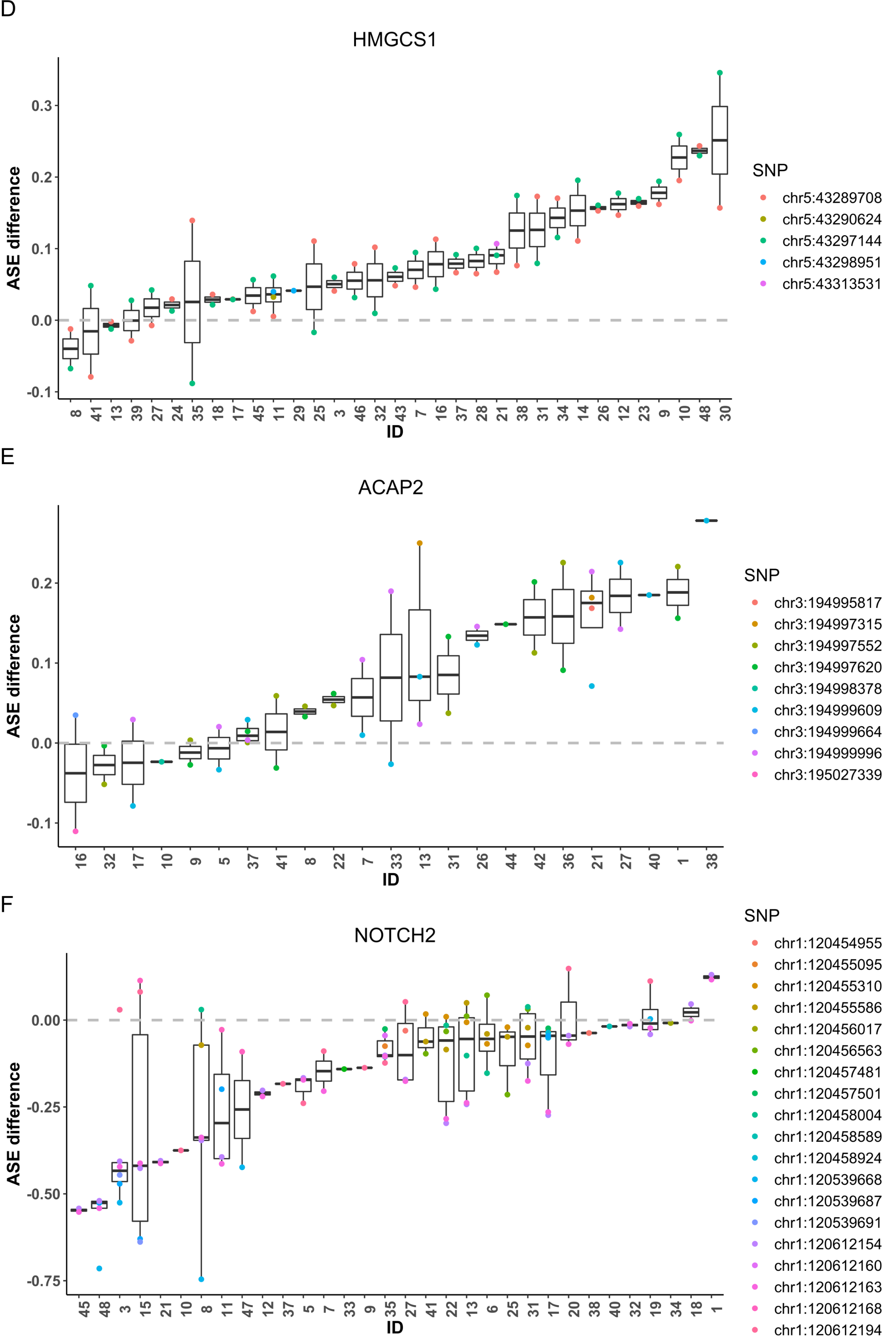

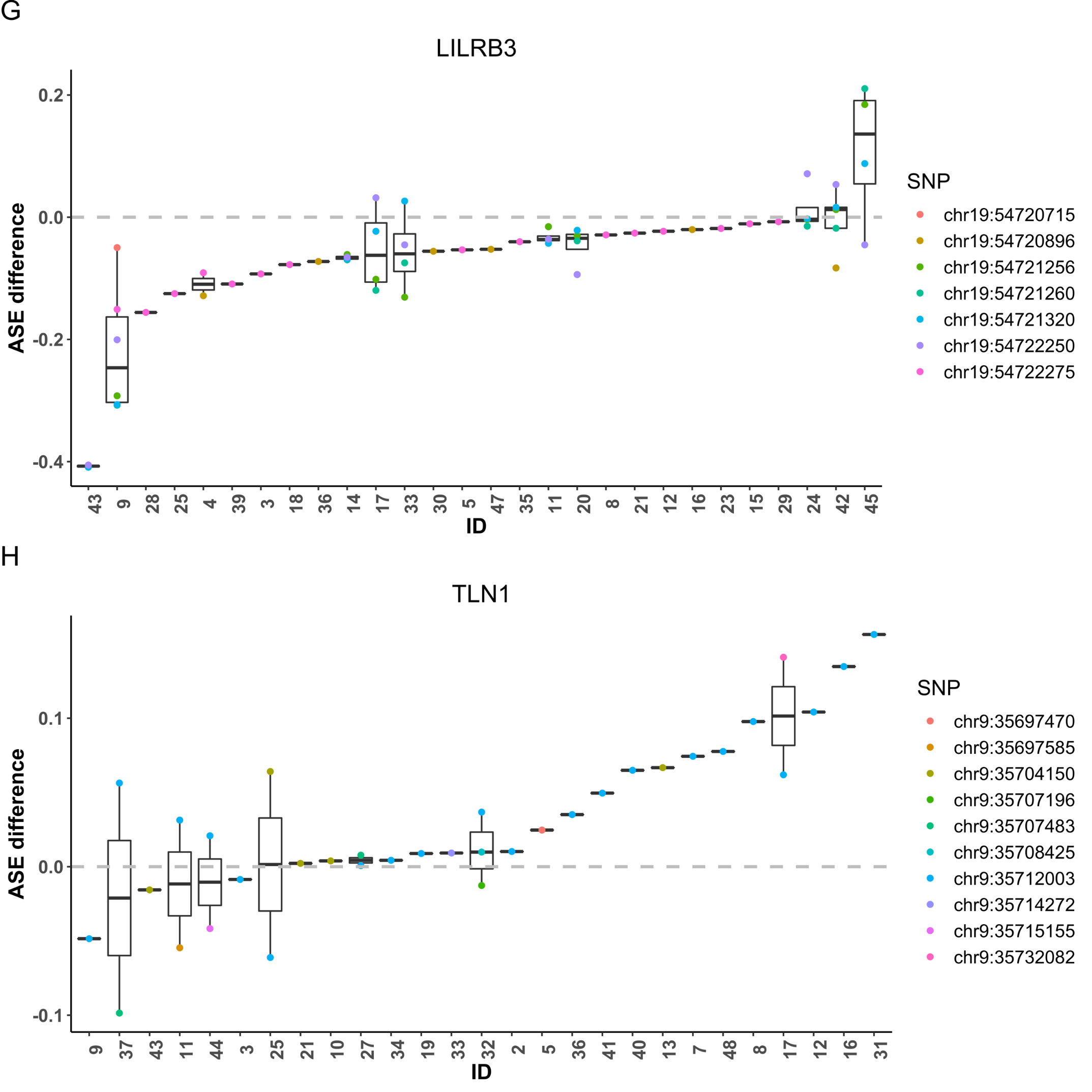

**Figure S3. SNP-level ASE difference between M0 and M1 macrophage samples for selected genes showing differential ASE in the macrophage RNA-seq dataset.** We selected eight genes to show their estimated SNP-level ASE difference across SNPs and individuals. The estimated ASE difference was obtained by calculating the major allele proportion difference between M1 and M0 samples after haplotype phase alignment. The individuals were sorted by median ASE difference across all SNPs.

**Supplementary Tables**

**Table S1. Significant ASE genes in M0 macrophage samples.** We detected 160 significant ASE genes (adjusted *P* < 0.05).

**Table S2. Significant ASE genes in M1 macrophage samples.** We detected 219 significant ASE genes (adjusted *P* < 0.05).

**Table S3. Significant differential ASE genes between M0 and M1 samples.** We detected 96 significant differential ASE genes (adjusted *P* < 0.05) between M0 and M1 macrophage samples.

**Table S4. Significant differential ASE genes between M0 and M1 macrophage samples that overlap with GWAS loci.** Among the 96 significant differential ASE genes, 52 genes overlap with GWAS results (*P* < 5x10-8) for cardiovascular disease, coronary artery disease, and acute coronary syndrome.

**Table S5. Subject demographics of the macrophage samples**

|  | Male | Female |
| --- | --- | --- |
| Sample number (N) | 24 | 24 |
| Age (median, range) | 35 (22-51) | 27 (21-50) |
| Race | Caucasian | Caucasian |
